# Opposing roles for iron transport systems in gallium tolerance in extraintestinal pathogenic *Escherichia coli*

**DOI:** 10.1101/2025.07.01.662525

**Authors:** Nagama Parveen, Seth Durrant, Michael A. Olson, Emily P. Shakespear, Trevor R. Jones, Eric Wilson, David L. Erickson

**Author notes:** Corresponding author: David Erickson, Department of Microbiology and Molecular Biology, Provo, UT, 84663.

## Abstract

Gallium is a promising antibacterial candidate because it displaces iron atoms inside bacterial cells but does not undergo redox cycling. It inhibits growth by disrupting essential iron-dependent processes. However, *Escherichia coli* are naturally less sensitive to gallium than many other bacteria, and the mechanisms that control gallium tolerance are not completely understood. We performed a genome-wide transposon sequencing (TnSeq) screen to identify genes important for the survival of an extraintestinal pathogenic *E. coli* isolate (M12) in gallium nitrate. The TnSeq results indicated that inactivation of enterobactin siderophore-related genes (*entS*, *fepD, fes, fepB*) enhances bacterial survival in gallium, while disrupting the ferric dicitrate transport system increases susceptibility. We validated these findings through targeted gene knockouts and gallium sensitivity experiments. Our findings suggest that enterobactin can complex with gallium for cellular uptake, but that the ferric citrate receptor FecA can discriminate between gallium citrate and iron citrate. Expression of *fecA* increased with gallium exposure, showing that gallium induces FecA-mediated iron uptake. Gallium also increased intracellular levels of manganese in the Δ*fecA* strain. Supplementation with iron or manganese restored growth of M12 Δ*fecA* in gallium, suggesting that gallium sensitivity is linked to both iron starvation and oxidative stress. As the ferric dicitrate transport system is an important virulence factor in several extraintestinal infection sites, our results suggest that targeting FecA may increase *E. coli* susceptibility to gallium while also suppressing virulence.

**Importance:** *Escherichia coli* extraintestinal infections that are resistant to traditional antibiotics are associated with more deaths than any other species. Gallium-based therapies may represent a non-antibiotic approach to treating extraintestinal pathogenic *E. coli* strains that affect both humans and animals. Our results are significant as they show that the enterobactin siderophore and the ferric dicitrate iron transport systems expressed by these bacteria have opposing roles in *E. coli* gallium sensitivity. These findings could be leveraged to enhance the efficacy of gallium therapeutics.

## Introduction

Extraintestinal pathogenic *Escherichia coli* (ExPEC) strains are responsible for many diseases in both humans and animals, including urinary tract infections, meningitis, pneumonia, and septicaemia (1, 2). ExPEC also infect mammary glands in lactating mammals (3, 4). For instance, dairy cattle frequently suffer from mastitis, which drastically lowers milk production and results in major economic losses. *E. coli* are responsible for 50-60% of severe acute mastitis cases which can often proceed to septicaemia and death (5). Multiple virulence factors have been implicated in mammary gland infection, such as activation of inflammation via pathogen-associated molecular patterns (6), complement resistance (7, 8), and adhesion and invasion of mammary epithelial cells (4, 9). Iron acquisition via the ferric dicitrate system is a critical mastitis virulence determinant (10, 11). The outer membrane receptor FecA senses and imports ferric citrate, which is then transferred through the periplasm and inner membrane by FecB, FecC, and FecD, powered by the ATPase FecE. Expression of the system is regulated by the extracytoplasmic sigma factor FecI and the anti-sigma factor FecR (12).

*E. coli* infections in livestock are commonly treated with antibiotics, which exerts selective pressure promoting resistance. Ready-to-eat animal products are frequently contaminated with *E. coli* strains resistant to a range of antibiotics, especially beta-lactams (13). Moreover, some *E. coli* strains isolated from meat also show resistance to last-resort antibiotics like carbapenems and colistin (14–16). Resistant *E. coli* strains are also often found in raw milk (17). Surging antimicrobial resistance in *E. coli* is a pressing global challenge affecting human and animal health.

Antimicrobial metals have been used for centuries to prevent bacterial growth and may represent one option to reduce antibiotic use. The potential of gallium to treat bacterial diseases has received increasing attention (18–22). Gallium and iron ions have similar nuclear radii, coordination chemistries, ionization potentials, electronegativity, and inclination to form ionic bonds. In bacteria treated with gallium, critical iron-containing proteins may be compromised as Ga^3+^ cannot be reduced under physiological conditions, unlike Fe^3+^ (23). Gallium compounds including gallium nitrate, gallium maltolate, gallium citrate, and gallium-protoporphyrin IX have efficacy against formidable antibiotic-resistant pathogens in animal models of disease (20, 24–26). Gallium compounds could be explored as part of a one-health approach to prevent food-borne illness or treat infected animals.

Although gallium compounds have antimicrobial activity in vitro and in vivo, many *Enterobacteriaciae* exhibit intrinsic resistance to gallium (27). The underlying genetic determinants of gallium resistance in *E. coli* remain poorly understood. Previous studies have identified genes that influence gallium resistance in a limited number of *E. coli* strains (28–33), including experimental evolution experiments, genetic screens, and gene expression analyses. Collectively, these studies implicate genes involved with iron transport systems as well as response to oxidative stress in gallium sensitivity. However, there is little overlap regarding the specific genes or pathways that are reported to be important and they may not be reflective of diverse clinical isolates belonging to different phylogroups. *E. coli* strains possess an arsenal of systems to maintain intracellular iron homeostasis, such as the ferric dicitrate transport system (Fec), siderophores including enterobactin, yersiniabactin, and aerobactin, and ferrous iron transporters EfeUOB or SitABCD (12, 34–36). Individual strains possess different combinations of these systems and may express them differently, which could impact their relative importance in surviving gallium stress.

We previously showed that prolific growth of the bovine mastitis clinical isolate M12 in mouse mammary gland infections depends on the ferric dicitrate transport system (10). This strain is also virulent in ascending urinary tract infection and sepsis models of disease (37). M12 also possesses genes for enterobactin, yersiniabactin, and SitABCD/EfeUOB transporters. In the present study, we used transposon sequencing to identify genes that impact fitness during exposure to gallium in strain M12 as an example of an ExPEC strain with zoonotic potential. In this study, we have uncovered opposing roles for the ferric dictrate and enterobactin iron acquisition systems in surviving gallium stress; the Fec system is induced by gallium stress and promotes growth, whereas the enterobactin system is detrimental. We also show an important protective role for manganese during gallium exposure.

## RESULTS

### Identification of genes that control gallium sensitivity in strain M12

There is a lack of information regarding genes required by ExPEC strains to survive in gallium nitrate. In preliminary experiments, we grew strain M12 (sequence type 69, phylogroup D) in Davis broth supplemented with various concentrations of gallium nitrate ranging from 1.56 µM – 1000 µM (data not shown) and found that 800 µM was slightly inhibitory but still allows growth. In order to identify genes that influence fitness of strain M12 in gallium nitrate, we utilized a transposon sequencing (TnSeq) approach. Starting with a library consisting of ∼400,000 unique Tn5 transposon insertion mutants [37], we pooled ten populations from the transposon mutant library and duplicate cultures were grown in Davis broth with or without 800 µM gallium nitrate. After an overnight incubation, the mutant populations were collected and sequenced in parallel from both conditions.

The output libraries were then mapped to the M12 genome and compared using DESeq2 (Figure 1). Genes whose disruption conferred a fitness change greater than 2 and a *P* value of 0.05 or lower are listed in Table 1. Genes that the TnSeq analysis indicated decrease fitness in gallium nitrate included the enterobactin exporter (*entS*), enterbactin esterase (*fes*), ferric enterobactin periplasmic (*fepB*) and inner membrane (*fepD*) transport proteins. The *ybeZ* gene encoding a predicted RNA helicase involved in 16S ribosomal RNA (38) was also detrimental to fitness based on the TnSeq results. We chose *entS*, *fepD*, and *ybeZ* for further investigation and created unmarked deletion mutants for each of these genes. As predicted by the TnSeq data, disruption of genes *entS, and fepD* in M12 led to better growth in gallium compared to the wild-type strain (Figure 2). Conversely, *ybeZ* deletion did not result in a statistically significant difference in growth in gallium (Figure 2).

**Figure 1.**
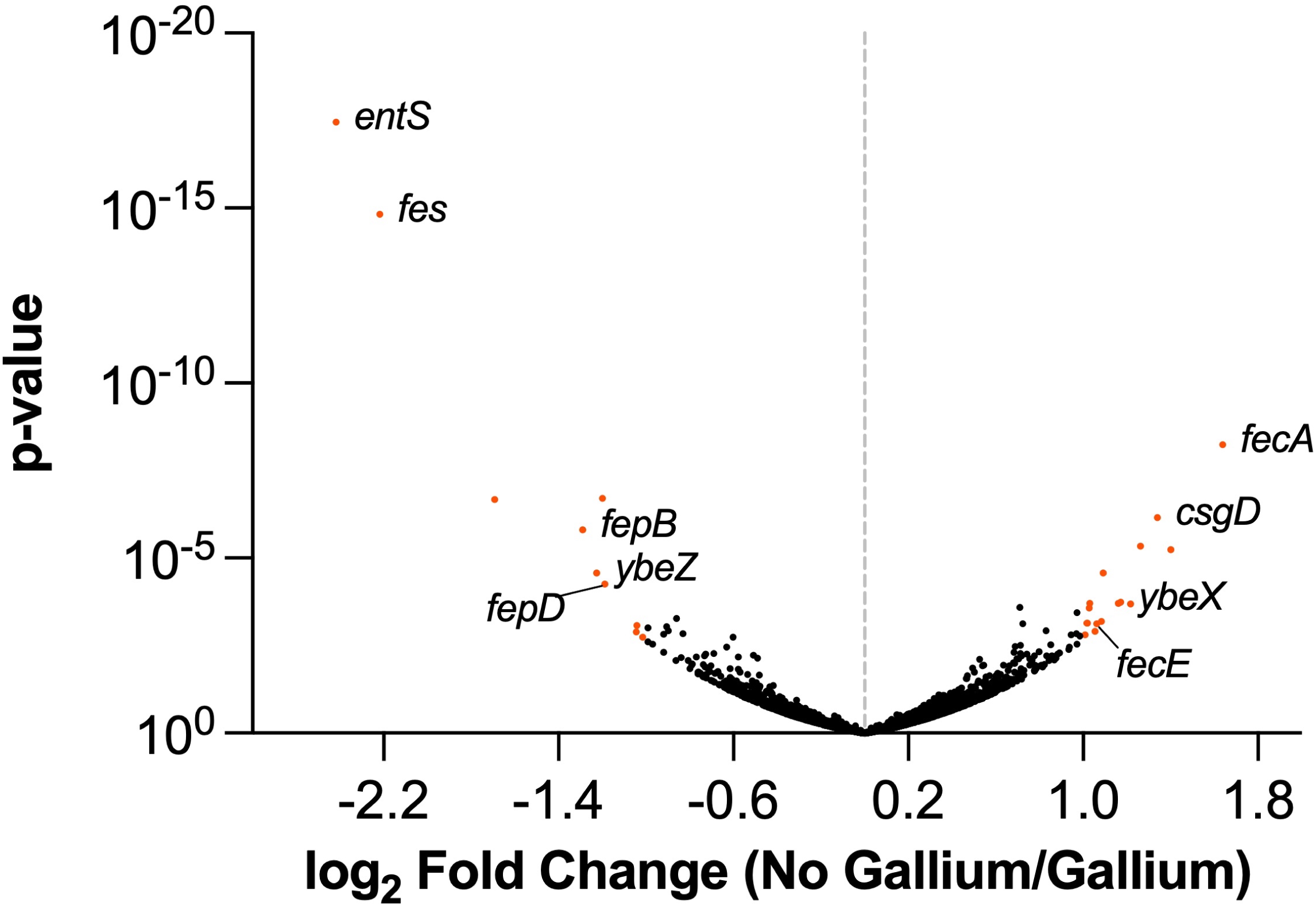
Genes controlling fitness of M12 during growth in gallium nitrate. Transposon sequencing was used to track mutants passaged in Davis broth or Davis broth supplemented with 800 µM gallium nitrate. Each dot on the volcano plot represents a gene with log_2_ fold change (reads mapped in no gallium/gallium) on the x-axis and the –log₁₀(p-value) on the y-axis. Genes with strong depletion (right) or enrichment (left) in the presence of gallium indicate loss or gain of fitness when disrupted by transposons, respectively. Orange dots indicate genes with a 2-fold change and p-value less than 0.05 as determined by DESeq2, and the names of genes investigated further are indicated.

**Figure 2.**
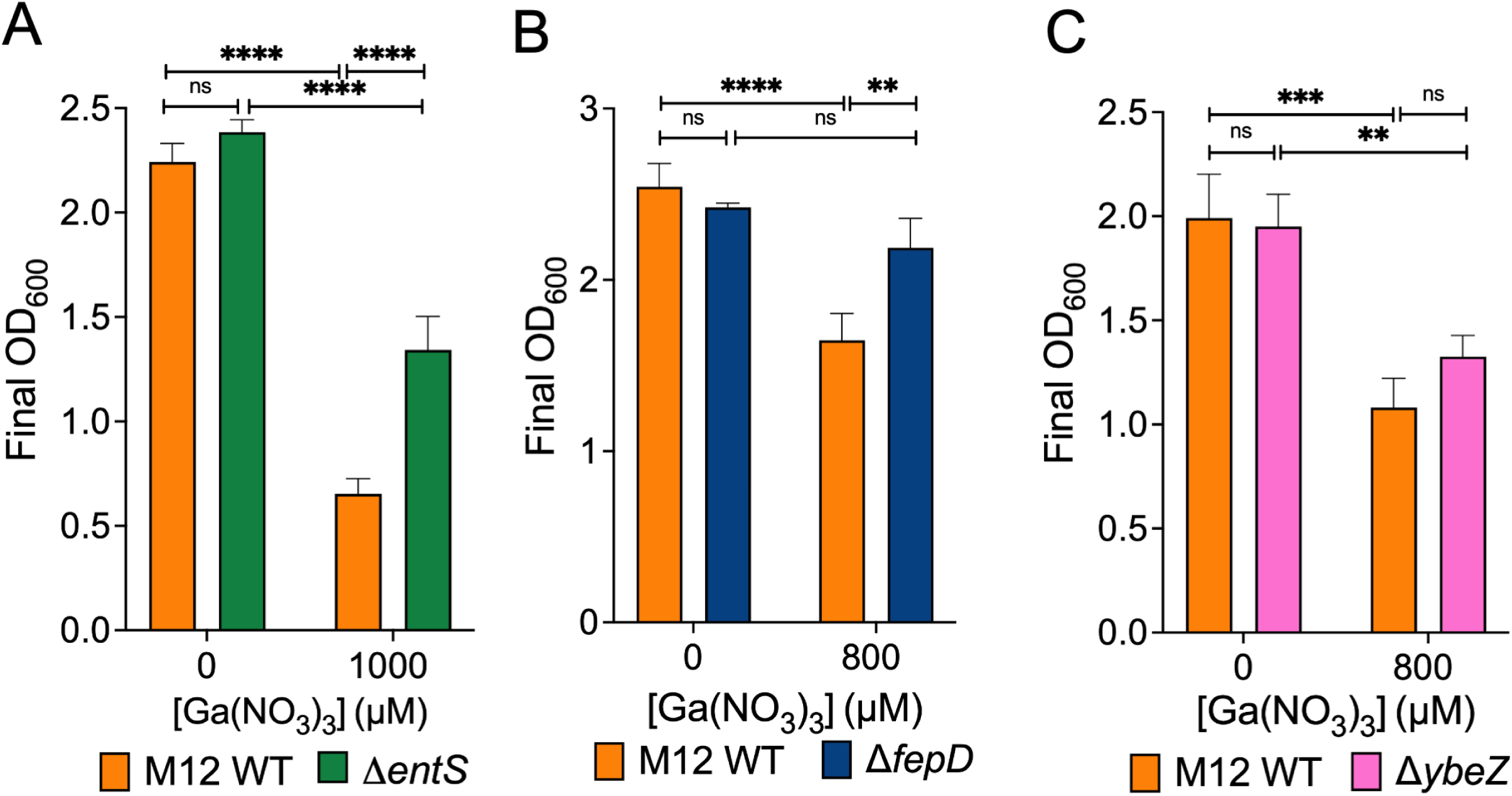
Enterobactin M12 mutants are more fit in gallium. Growth of the M12 wild-type strain was compared with (A) Δ*entS* (B) Δ*fepD* or (C) Δ*ybeZ* mutant strains in the presence or absence of gallium nitrate. * = p-value < 0.05, ** = p-value < 0.01 by two-way ANOVA. Shown are the mean ± standard deviation of triplicate samples from a single representative experiment performed at least twice.

**Table 1.** Top genes affecting fitness of *E. coli* M12 during growth in gallium nitrate.

| Name | Log2 Ratio | p-value |
| --- | --- | --- |
| Enterobactin exporter EntS | -2.42 | 3.50E-18 |
| Enterochelin esterase Fes | -2.22 | 1.47E-15 |
| Exodeoxyribonuclease 7 small subunit XseB | -1.69 | 2.08E-07 |
| Ferrienterobactin-binding periplasmic protein FepB | -1.29 | 1.57E-06 |
| PhoH-like protein YbeZ | -1.23 | 2.64E-05 |
| 5-keto-D-gluconate 5-reductase IdnO | -1.20 | 1.93E-07 |
| Ferric enterobactin transport system permease protein FepD | -1.18 | 5.51E-05 |
| hypothetical protein CDS | -1.04 | 0.0013 |
| hypothetical protein CDS | -1.04 | 0.00084 |
| YihD | -1.02 | 0.0018 |
| Fe(3+) dicitrate transport protein FecA | 1.64 | 5.69E-09 |
| Sugar phosphatase YbiV | 1.40 | 5.74E-06 |
| CsgBAC operon transcriptional regulatory protein CsgD | 1.34 | 7.02E-07 |
| UvrABC system protein B | 1.26 | 4.54E-06 |
| hypothetical protein CDS | 1.22 | 0.00020 |
| YbeX/CorC ribosome assembly protein | 1.17 | 0.00018 |
| D-alanyl-D-alanine carboxypeptidase DacA | 1.16 | 0.00019 |
| putative multidrug resistance ABC transporter ATP-binding/permease protein MdlB | 1.09 | 2.62E-05 |
| Anaerobic dimethyl sulfoxide reductase chain C | 1.08 | 0.00063 |
| Fe(3+) dicitrate transport ATP-binding protein FecE | 1.06 | 0.00076 |
| putative protein YceO CDS | 1.06 | 0.00125 |
| 50S ribosomal protein L31 type B | 1.03 | 0.00019 |
| hypothetical protein CDS | 1.03 | 0.00027 |
| Transcriptional regulatory protein CpxR | 1.02 | 0.00072 |
| hypothetical protein | 1.02 | 0.00072 |
| Multidrug efflux pump subunit AcrA | 1.01 | 0.00154 |

Genes that TnSeq predicted would enhance gallium resistance included those with clear roles in surviving antimicrobial stress, including the efflux genes *mdlA* (39) and *acrA* (40), envelope stress transcriptional regulator *cpxR* (41), and the cell wall carboxypeptidase *dacA* (42). Transposon sequencing indicated that the ribosome assembly protein *ybeX* (43), encoded in the same operon as *ybeZ,* as well as the *csgD* transcriptional regulator gene (44) are needed for gallium resistance. Notably, the ferric dicitrate transporter *fecA* as well as the *fecE* gene encoding the ATPase for the system were also identified in the TnSeq screen. We chose to create mutants lacking *csgD*, *ybeX*, and *fecA* for further investigation. Deletion of *csgD* did not reduce growth in gallium compared to the wild-type strain (Figure 3A). Disruption of *ybeX* proved modestly detrimental to M12 growth in gallium (Figure 3B). A much more pronounced defect was seen when *fecA* was deleted (Figure 3C). Wild-type levels of growth were restored to the *fecA* mutant complemented with a plasmid encoding the *fecABCDE* operon (pMO2). Growth of the wild type, Δ*fecA* mutant, and complemented strains were measured over a range of gallium nitrate concentrations to determine the sensitivity of the mutant more precisely. The M12 Δ*fecA* mutant grew as well as the wild-type strain at 200 µM gallium nitrate (Figure 3D).

**Figure 3.**
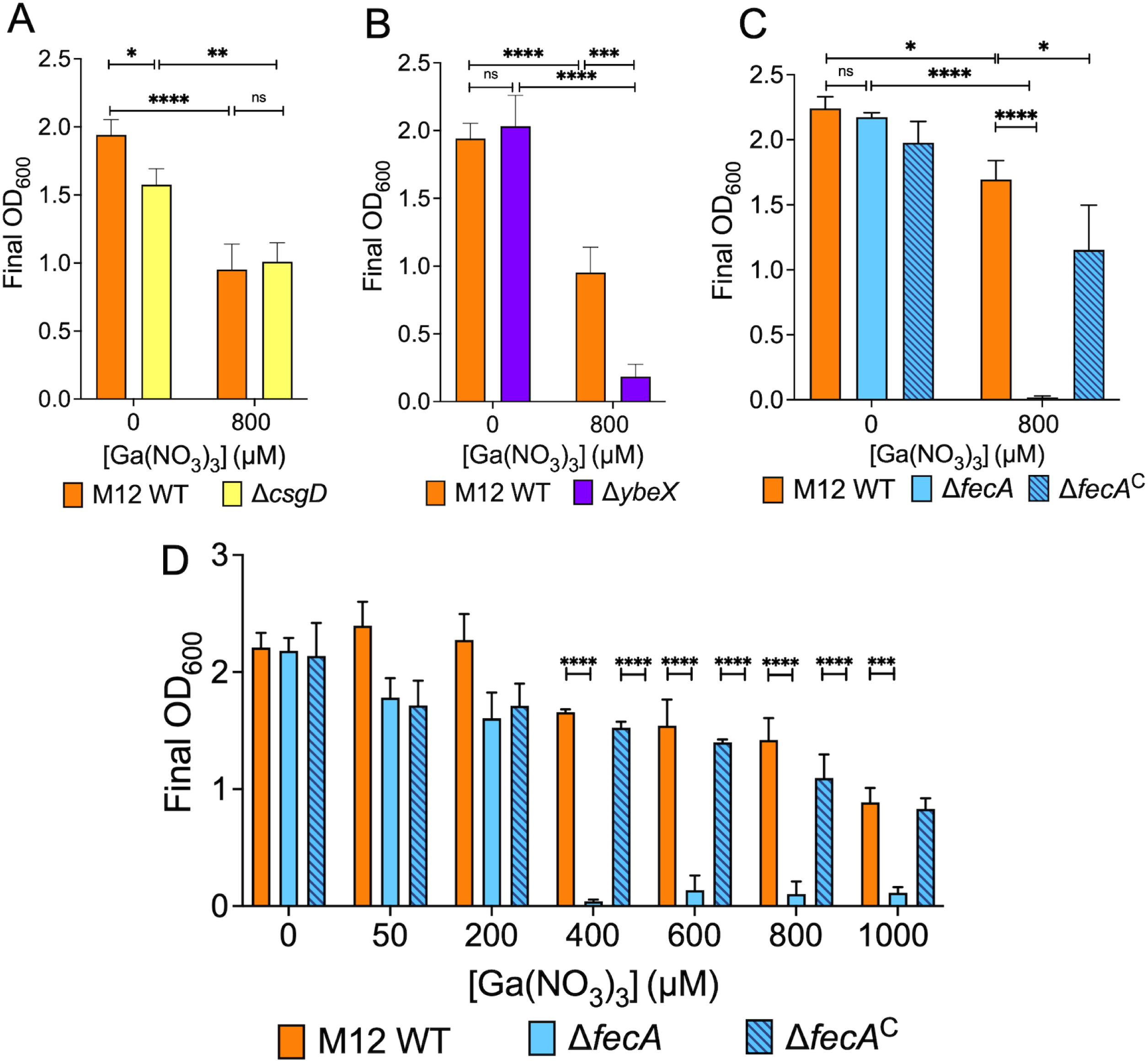
The ferric dicitrate transporter *fecA* enhances M12 fitness in gallium. (A-C) Growth yields of M12 WT were compared with mutants Δ*csgD* (A), Δ*ybeX* (B) or Δ*fecA* and the Δ*fecA*^C^ complemented mutant containing plasmid pMO2 (C) in Davis broth with or without 800 µM gallium. (D) Growth of wild-type (M12), mutant (Δ*fecA*), and Δ*fecA*^C^ in gallium concentrations ranging from 0 µM – 1000 µM. * = p-value < 0.05, ** = p-value < 0.01, *** = p-value < 0.001, **** = p-value < 0.0001 by two-way ANOVA. Shown are the mean ± standard deviation of triplicate samples from a single representative experiment.

To determine whether *fecA* is required for other mastitis-associated strains to grow in the presence of gallium, we removed *fecA* from three additional clinical isolates (M34, M44, and M66) and exposed them to 800 µM gallium nitrate. Clear growth defects were observed in strains lacking *fecA* in the presence of gallium for each (Figure 4A-C), confirming its importance in multiple mastitis strains. However, it is evident that M12 Δ*fecA* is more sensitive than the M34, M44, and M66 Δ*fecA* mutants. Additionally, the wild type M44 and M66 strains were not inhibited by 800 µM gallium nitrate. We also observed a slight growth defect of the *fecA* mutants of strains M34, M44 and M66 in Davis broth without gallium. Not all mastitis-associated *E. coli* strains contain the *fecABCDE* operon, and thus may be more sensitive to gallium nitrate than those that do. Strain M3 lacks *fecA* and it has a growth defect at 500 µM, but introducing plasmid pMO2 encoding the *fecABCDE* operon significantly enhanced growth in gallium nitrate (Figure 4D).

**Figure 4.**
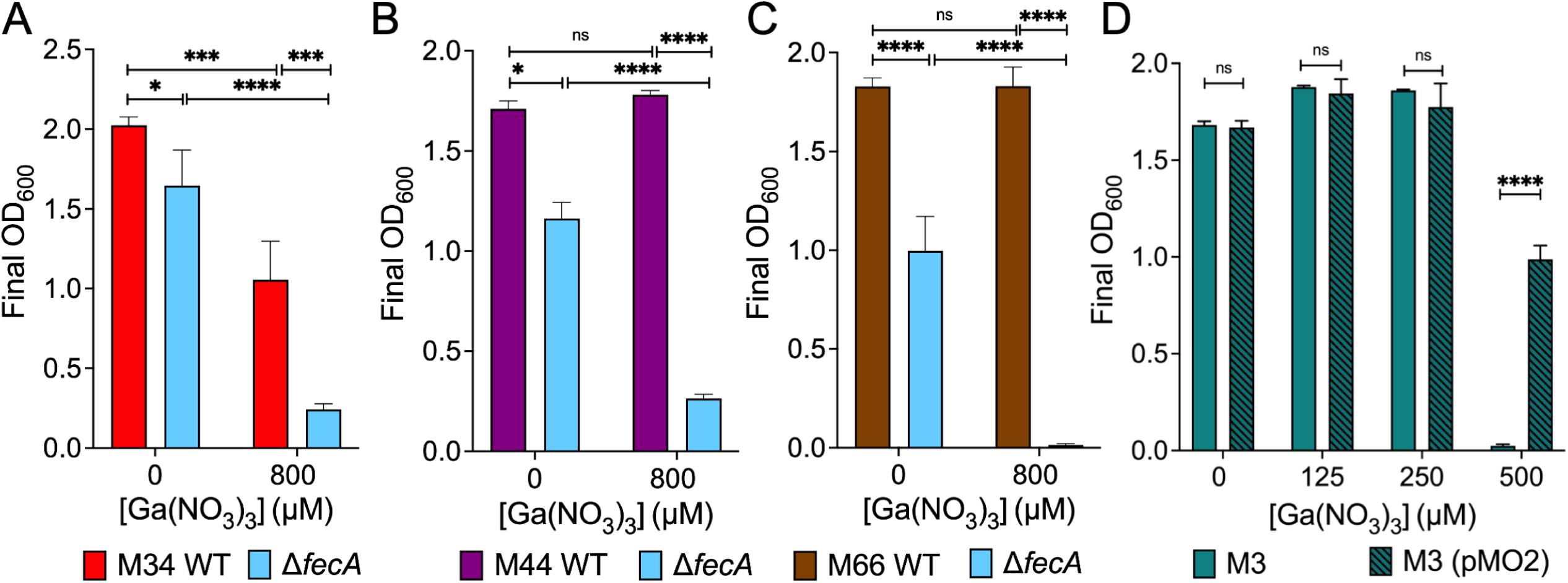
The ferric dicitrate system increases gallium tolerance in multiple mastitis-associate *E. coli* strains. (A-C) Growth of mastitis-associated strains M34, M44, and M66 and their Δ*fecA* mutants in Davis broth with or without 800 µM gallium. (D) Growth of mastitis-associated strain M3, which lacks the *fecIR* and *fecABCDE* genes, and M3 (pMO2) was compared in different concentrations of gallium nitrate as indicated. * = p-value < 0.05, ** = p-value < 0.01, *** = p-value < 0.001, **** = p-value < 0.0001 by two-way ANOVA. Shown are the mean ± standard deviation of triplicate samples from a single representative experiment.

### Supplementation with iron or manganese overcomes the growth defect of *fecA* mutant

The M12 Δ*fecA* mutant may be less fit in gallium because it is unable to acquire enough iron to outcompete the gallium in the media. To test this, we measured bacterial growth of the wild type and mutant strains in Davis broth supplemented with ferrous sulphate, ferric chloride, and manganese sulphate in the presence of 800 µM gallium. Ferrous sulfate completely rescued the growth of M12 Δ*fecA* at 6.25 µM (Figure 5A), whereas 50 µM ferric chloride was required to rescue growth (Figure 5B). A relatively small amount of supplementary iron was required to restore growth of the Δ*fecA* mutant compared to the gallium in the media (6.25 vs 800 µM), suggesting that iron is much more readily taken up than gallium, or that gallium competes poorly for metal binding sites in critical enzymes. Interestingly, manganese was more potent than ferrous iron (Figure 5C), as only 1.56 µM was required to restore growth.

**Figure 5.**
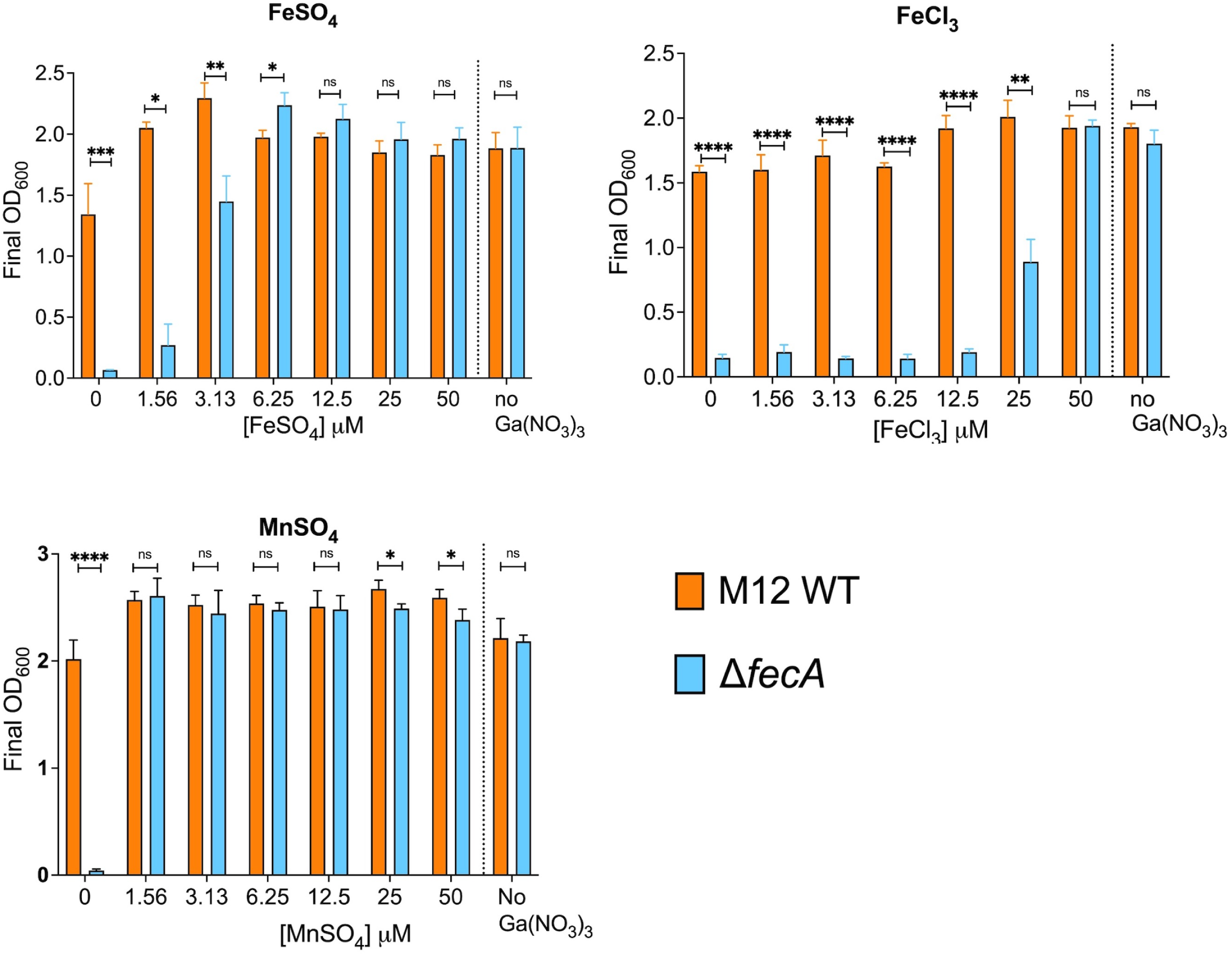
Gallium-mediated growth inhibition of Δ*fecA* is alleviated by co-supplementation with manganese or iron. Growth yields of M12 wild-type or Δ*fecA* in Davis broth with 800 µM gallium supplemented with (A) FeSO_4_, (B) FeCl_3_ or (C) MnSO_4_. * = p-value < 0.05, ** = p-value < 0.01, *** = p-value < 0.001, **** = p-value < 0.0001 by two-way ANOVA. The mean ± standard deviation of triplicate samples from a single representative experiment are shown.

### Loss of FecA reduces intracellular iron and increases manganese uptake in the presence of gallium

To understand how loss of *fecA* affects metal homeostasis during gallium stress, we quantified intracellular accumulation of iron, zinc, gallium, and manganese in the wild-type and the mutant strains using ICP-MS (Figure 6). In the wild-type strain, the presence of 200 µM gallium in the media resulted in slightly less intracellular iron, manganese, and zinc, although this difference was only statistically significant for zinc. Conversely, the iron content of M12 Δ*fecA* mutant was reduced to roughly half that of wild-type M12 when grown in gallium. Equivalent iron levels were observed in the M12 wild type and Δ*fecA* mutant in the absence of gallium. Conversely, intracellular gallium levels were slightly higher in M12 Δ*fecA* mutant than the wild-type strain grown in the presence of gallium, demonstrating that FecA is not the major route for gallium entry. Growth in gallium strongly elevated manganese levels the Δ*fecA* mutant compared with the wild type strain. When gallium was present in the media, intracellular zinc was significantly decreased in both wild-type and Δ*fecA*, which was even lower in the Δ*fecA* mutant compared with the wild-type strain.

**Figure 6.**
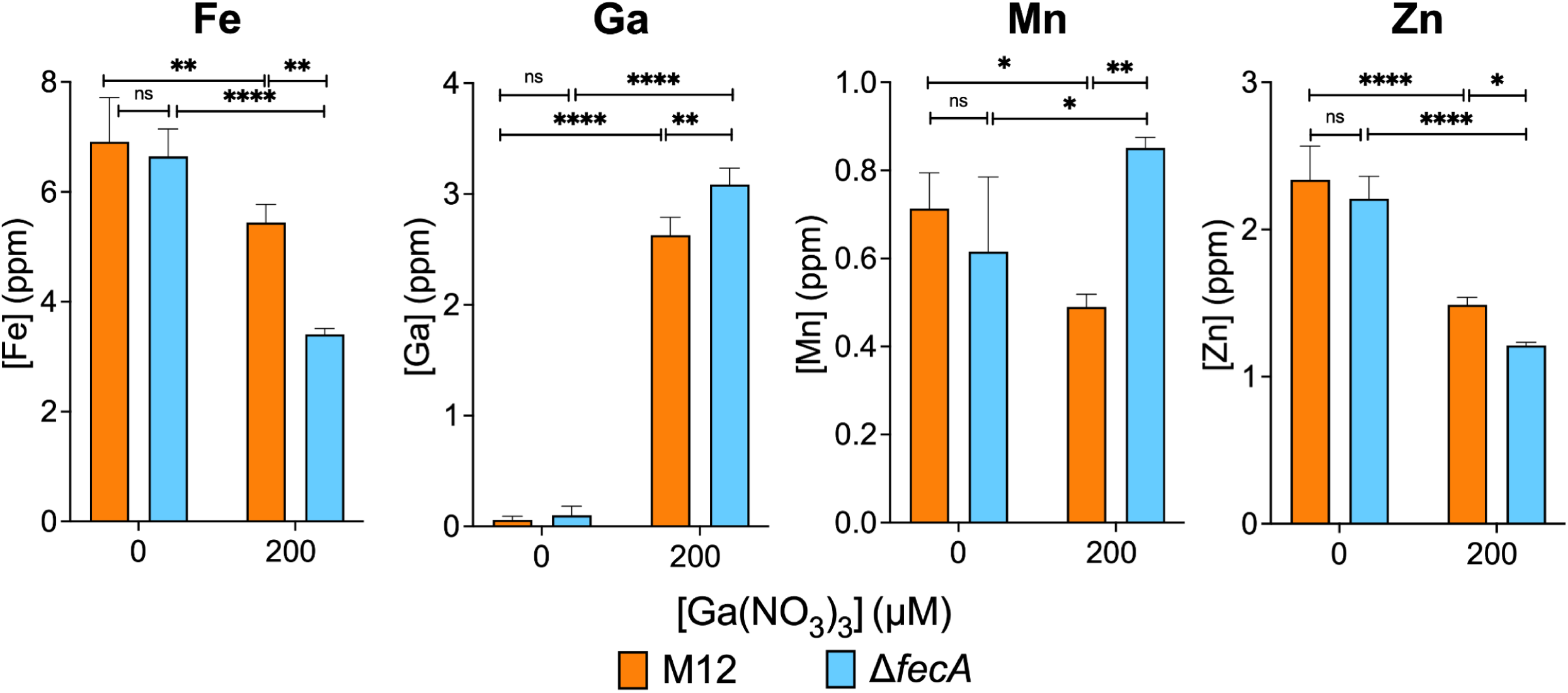
Intracellular accumulation of metals is affected by growth in gallium and loss of FecA. M12 WT and Δ*fecA* were grown in the presence or absence of 200 µM Ga(NO_3_)_3_ and washed bacterial cells analyzed by ICP-MS for iron, gallium, manganese, and zinc. * = p-value < 0.05, ** = p-value < 0.01, *** = p-value < 0.001) by two-way ANOVA. Shown are the mean ± standard deviation (error bars) of triplicate samples from a single experiment performed three times.

### *fecA* expression is induced by gallium

Expression of *fecA* is active when intracellular iron levels are low and by the presence of ferric citrate in the environment. We wanted to determine whether gallium would affect *fecA* expression by complexing with citrate to induce signalling. We created a transcriptional reporter wherein the *lacZ* gene was placed under the control of the previously defined *fecA* promoter (45). The same plasmid without the *fecA* promoter was used as a control. As shown in Figure 7, β-galactosidase activity from the *fecA::lacZ* plasmid was increased by addition of gallium to the growth media in a dose-responsive manner, unlike the empty vector or the untransformed parental M12 strain.

**Figure 7.**
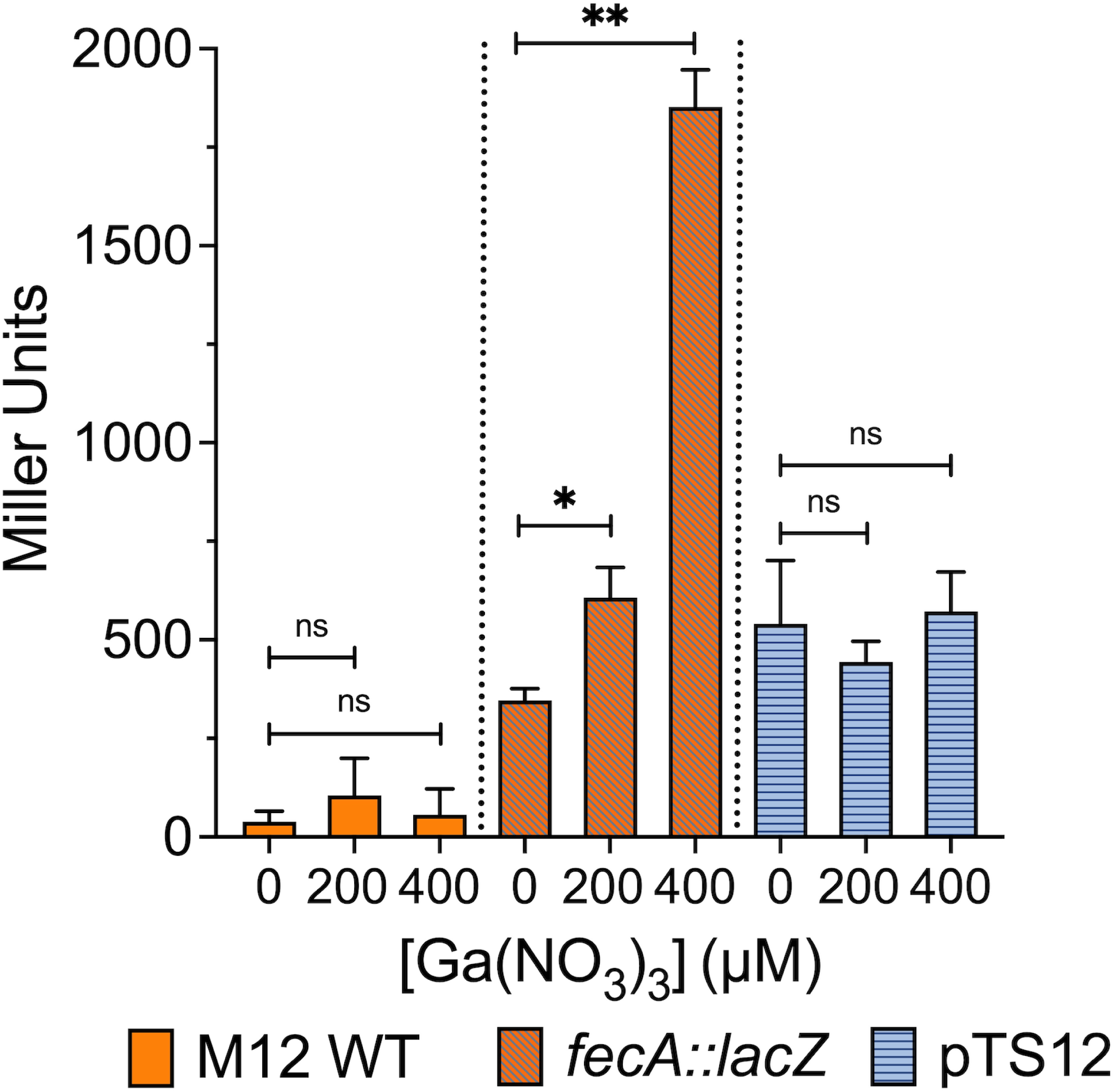
Gallium upregulates the expression of *fecA* promoter in M12. ß-galactosidase reporter assay was used to measure *fecA* expression in response to Ga(NO_3_)_3_ in the media by Miller assay. The untransformed parental M12 strain as well as the promoterless vector only strain were used as controls. The 200 and 400 µM values were compared with the no gallium control by one-way ANOVA, * = p-value < 0.05, ** = p-value < 0.01. Shown are the mean ± standard deviation (error bars) of triplicate samples from a single representative experiment performed at least twice.

### Loss of FecA also increases sensitivity to cobalt

To determine whether the M12 Δ*fecA* mutant is also more sensitive to other antimicrobial metals, we measured the MIC of strains M12 and M34 and their respective Δ*fecA* mutants to silver nitrate, copper sulphate, and cobalt chloride. The MICs for silver nitrate and copper sulphate were the same for the wild type and Δ*fecA* mutant strains (Table 2). However, the mutants showed two-fold lower MIC for cobalt chloride both M12 and M34 backgrounds, suggesting that iron citrate import helps counter stress induced by cobalt as well as gallium.

**Table 2.** MIC of antimicrobial metals for mastitis-associated *E. coli* and Δ*fecA* mutants.

| Strains | CuSO<sub>4</sub> (μM) | AgNO<sub>3</sub> (μM) | CoCl<sub>2</sub> (μM) |
| --- | --- | --- | --- |
| M12 WT | 500 | 50 | 313 |
| M12 $\Delta fecA$ | 500 | 50 | 157 |
| M34 WT | 1000 | 50 | 1250 |
| M34 $\Delta fecA$ | 1000 | 50 | 625 |

## Discussion

The iron-mimicking properties of gallium make it toxic to many bacteria. However, *E. coli* exhibit natural resistance. Therefore, this work aimed to understand genes possessed by mastitis-associated *E. coli* that limit gallium uptake or mitigate its toxic effects. We identified several genes, particularly those involved in iron acquisition, that significantly influenced M12 fitness in gallium (Figure 1). Transposon insertions in enterobactin siderophore genes like *entS*, and *fepD* allowed M12 to survive in gallium. Conversely, insertions in the ferric dicitrate receptor gene *fecA* resulted in greater sensitivity to gallium, indicating a protective role of the gene. This surprising result was verified with deletion mutants of enterobactin secretion (*entS*) and receptor genes (*fepD*) (Figure 2) as well as the ferric dicitrate receptor (*fecA*) in multiple strains (Figures 3 and 4). The most straightforward interpretation of these findings is that enterobactin binds gallium and gallium-enterobactin complexes are imported. Thus, eliminating enterobactin receptor or transport genes prevents gallium entry, and eliminating the enterobactin exporter is especially protective since it concentrates the siderophore in the cytosol where it can neutralize intracellular gallium. Conversely, although citrate complexes with either iron or gallium, we propose that the ferric dicitrate receptor FecA transports only iron. Gallium citrate can induce signalling leading to *fecA* expression (Figure 7). However, the FecA and/or the periplasmic/inner membrane components (FecB, FecC, FecD) are selective for iron import (Figure 6). Thus, deletion of these genes reduces intracellular iron that is necessary to compete with more abundant gallium that enters by other means (such as the enterobactin pathway).

To our knowledge, no prior studies have reported such opposing roles of *E. coli* iron transport systems in the context of gallium susceptibility. In an adaptive evolution study (28) wherein *E. coli* developed resistance to gallium under selective pressure, Graves et al. identified polymorphisms in the intergenic regions of *fepD* and *entS* in the gallium-resistant population. The effects of these intergenic SNPs on enterobactin gene expression or gallium resistance were not determined. We have shown that mutations in the enterobactin pathway increase gallium resistance, but we did not detect a role for yersiniabactin, which is also made by strain M12. Pyoverdine (46) or pyochelin siderophore (47) mutants *of Pseudomonas aeruginosa* are also more resistant to gallium. Thus, siderophore-mediated gallium uptake appears to be common in Gram-negative bacteria.

Graves et al. also noted the selection of two polymorphisms in *fecA* in gallium-resistant *E. coli* populations, but they did not verify the impact of these changes on gallium resistance. Both mutations affect the citrate binding pocket of FecA (48, 49). The first mutation resulted in a glycine to serine substitution at position 400, directly adjacent to the citrate binding site. It is possible that the polar side chain of serine hinders ferric citrate binding, but this has not been tested. It is worth noting that a different bacterial iron-binding protein, cytochrome c_2_ of *Rhodobacter capsulatus*, also has a glycine directly adjacent to the ligand binding site (50). When this residue was changed to serine, the protein maintained its stability and in vivo function, but the network of hydrogen bonds in the protein were altered. The second mutation changed asparagine to lysine at residue 169, which is part of a helix that faces away from the citrate pocket. The larger, charged R-group of lysine might push the helix into the binding pocket and change affinity for the ligand. Graves et al. hypothesized that these *fecA* mutations impair its ability to transport ferric citrate, and by extension gallium citrate, leading to higher gallium tolerance. Our results imply that FecA is not a significant contributor to gallium citrate import, but instead imports iron even when gallium citrate is abundant. Support for this argument also comes from direct measurement of gallium internalized by *P. aeruginosa*, which possesses a ferric dicitrate receptor. When exposed to equal concentrations of either iron or gallium citrate, the bacteria internalize much more iron than gallium (51), and gallium citrate is not imported more efficiently than gallium nitrate by these bacteria (47), consistent with strong selectivity for iron citrate of the Fec system.

Deleting *fecA* not only eliminates transport of ferric citrate but also disrupts the FecA-FecR-FecI signaling cascade. FecI upregulates the *fecIR* and *fecABCDE* operons (52), but might also recruit RNA polymerase to other locations. Additionally, FecI induction likely affects expression of a wide variety of genes indirectly through competition with other sigma factors (53, 54). Gallium sensitivity of Δ*fecA* mutants might be at least partially due to these other putative gene expression changes. However, expression of the *fecABCDE* operon alone in strain M3 increased its gallium resistance (Figure 4). This plasmid does not include the *fecI* or *fecR* genes. Therefore, an alternative explanation for the findings of Graves et al. might be that the subtle changes within *fecA* in gallium-resistant populations do not eliminate protein function but further enhance its iron selectivity over gallium, or enable a stronger signalling response leading to more FecA production. Compensatory mutations that overcome the fitness cost of reduced *fecA* function are also possible.

Most iron acquisition genes are co-ordinately regulated by Fur, including the ferric dicitrate, yersiniabactin, and enterobactin systems. In a strain possessing only enterobactin and the *fec* genes, inactivating the enterobactin pathway increases expression of the *fec* genes (55). This could also partially explain why the M12 *entS* mutant survives better in gallium (Figure 2). In this context, *fecA* might be upregulated to supply needed iron, and the trapped enterobactin inside the cell chelates gallium, preventing it from disrupting iron homeostasis. In contrast, when *fecA* is absent, the cell may overexpress the enterobactin system, thereby importing more gallium and exacerbating its effects. Whether strains such as M12 that contain multiple iron-acquisition systems including yersiniabactin upregulate enterobactin production when the Fec system is disrupted should be investigated.

We hypothesized that the M12 Δ*fecA* mutant sensitivity to gallium was due to failure to acquire enough iron to outcompete gallium. The M12 Δ*fecA* needed only 6.25 µM or 1.56 µM of supplemental ferrous iron or manganese, respectively, to completely restore the growth in the presence of 800 µM gallium, but a significantly higher (50 µM) concentration of ferric iron (Figure 5). This may be due to the low solubility of ferric iron under aerobic conditions, or to the absence of *fecA*, a key transporter for ferric iron. Manganese was more effective than iron in rescuing the growth of M12 Δ*fecA.* This likely stems from its role in antioxidant enzymes like Mn-SOD or ribonucleotide reductase NrdEF in low iron conditions (56, 57). Manganese supplementation also restored gallium resistance in an *E. coli* strain lacking all iron/manganese transporters (58). We found that loss of *fecA* and exposure to gallium resulted in increased intracellular manganese (Figure 6), implying that iron shortage and oxidative stress may be compensated by an active increase in manganese uptake. This may stem from upregulation of the *mntH* gene during iron starvation, which codes for a transport system that prefers to uptake Mn²⁺ but can also import Fe²⁺ (55).

The intracellular iron level in the M12 Δ*fecA* mutant was approximately half that of the wild type strain when 200 µM gallium was present (Figure 6). In the wild-type strain, iron levels were only slightly lower when gallium was present and this was not statistically significant. This suggests that the wild-type strain may upregulate iron transport genes when gallium is present, particularly the *fecABCDE* genes. We observed a clear upregulation of the *fecA* promoter in Davis broth supplemented with gallium (Figure 7), consistent with a cellular response to perceived iron limitation and ferric citrate (or gallium citrate) availability. A recent transcriptomic study also reported upregulated Fe^2+^ and Fe^3+^ import systems under gallium nitrate stress, including *fecABCDE* in *E. coli* (32). Taudte et al. (58) also found that intracellular iron levels were unaffected by gallium in the growth media. We detected slightly more intracellular gallium in the Δ*fecA* mutant than the wild-type strain (Figure 6). Collectively, these results indicate that gallium probably does not enter by FecA but can enter through other iron transport systems. These transporters are much more efficient for iron than gallium, and additional transporters for gallium remain to be identified.

Gallium in the media reduced intracellular zinc, and the levels were lower in M12 Δ*fecA* than the wild-type strain (Figure 6D). Gallium might interfere with uptake via zinc transporters such as ZupT. ZupT is known to bind zinc, cadmium, and iron with high affinity (59); it is possible that gallium competes with iron for uptake through this pathway.

Loss of *fecA* also impacted sensitivity to cobalt, but not copper or silver (Table 2). Cobalt can be incorporated during the synthesis of iron-sulfur [Fe-S] clusters in vital metabolic proteins, causing the proteins to become inactive (60). Strains deficient in the machinery involved in the assembly of [Fe-S] cluster are hypersensitive to cobalt, and cobalt treatment increases the expression of iron uptake genes (60). This model is consistent with our observation of elevated sensitivity of the Δ*fecA* mutant, which might be unable to fully replenish the [Fe-S] clusters damaged by cobalt. Future work should include testing whether cobalt and gallium may be synergistic, especially when FecA is disrupted.

In addition to the iron-associated genes, the TnSeq results identified two other genes (*ybeX* and *ybeZ*) found in the same operon that were predicted to influence M12 survival in gallium. YbeX is associated with ribosome metabolism (43), and the absence of this gene was confirmed to increase M12 sensitivity to gallium (Figure 3). On the other hand, YbeZ is a putative RNA helicase that has phosphatase activity (38), the absence of which did not detectably affect M12 growth in gallium (Figure 2). YbeX is known to play a role in temperature and antibiotic stress responses (43). The absence of *ybeX* potentially amplifies ROS-mediated damage to ribosomes in the presence of gallium.

Enhancing the antimicrobial potency of gallium may be a promising approach to combat drug resistance, including in agricultural settings. This will require a complete understanding of the specific proteins that are targeted by gallium, as well as the physiological responses that govern sensitivity. Our study contributes to this understanding by highlighting the important role *fecA* plays in maintaining iron homeostasis, which helps mitigate gallium toxicity. Moreover, we have uncovered the protective role of manganese under gallium stress by helping the cell survive under these conditions. Our results might inform future strategies to make gallium more effective against *E. coli*. Specifically, although gallium citrate is not likely to be effective, complexing gallium to enterobactin might be. Simultaneously interfering with the FecA and MntH proteins with antibodies, chemicals, or aptamers could make an enterobactin-gallium conjugate even more potent. Given the important role of the Fec and MntH systems in ExPEC urinary tract infections (55), sepsis (61), and mastitis (10), this strategy might have broad applicability.

## Materials and Methods

### Bacterial strains and conditions

*E. coli* strains M3, M12, M34, M44, and M66 were previously isolated from milk samples from cattle with clinical mastitis [1, 2]. For routine growth, strains were streaked on Luria-Burtani agar from freezer stocks and grown at 37°C. Liquid cultures were grown in Davis broth (62) supplemented with 0.1% (w/v) casamino acids. Under conditions where bacteria were grown in the presence of gallium, a fresh 1 M stock of gallium nitrate hydrate (Sigma-Aldrich) was prepared in 70% ethanol, which was diluted as needed to reach the desired concentration in Davis broth. When required, growth media was supplemented with kanamycin (50 µg/ml) or ampicillin (100 µg/ml).

### Transposon Sequencing

To prepare DNA for transposon sequencing, ten vials of an M12 transposon mutant library (10) were pooled and grown in Davis Broth overnight. The following day, duplicate subcultures were prepared in Davis broth under two conditions: one without Ga(NO_3_)_3_ and the other with 800 µM (Ga(NO_3_)_3_). These cultures were grown overnight, and total DNA was isolated from each using gBAC mini genomic DNA kit (IBI Scientific). The transposon insertion junctions were amplified and prepared for sequencing as previously described (10). Illumina 100 bp reads were generated on a NextSeq 2000 P1 Illumina reads (single ends) at SeqCenter (Pittsburgh, PA). A closed, high quality genome sequence of strain M12 was generated by mapping previously generated Illumina short reads (10) to Oxford Nanopore long read assembly generated by Plasmidsaurus (Louisville, KY). The sequences corresponding to each sample were separated using their barcodes, and mapped to the M12 genome using the STAR aligner (63). For control samples, 2,066,541 and 1,812,044 reads mapped to the M12 genome in replicates one and two, respectively. In the gallium treated samples, 1,383,488 and 1,336,748 reads were mapped for the first and second replicates, respectively. Raw count tables for each gene were analysed using the DESeq2 package in Geneious (Biomatters). The fold change values were log_2_ transformed, and a *P* value was calculated using a negative binomial distribution performed by DESeq2. Candidate genes contributing to fitness in gallium were identified by a log_2_-fold change >2 or <-2 and a *P* value of 0.05 or less.

### Gene deletion

Unmarked deletions were created by allelic exchange using plasmids pFOK (64) or pAX1 (65). Approximately 500 bp upstream and downstream regions of the gene of interest were PCR amplified with Q5 high fidelity mix (NEB) using primers listed in Supplementary Table 1. For cloning, the vector was linearized with EcoR1 (pFOK) or SalI and AvrII (pAX1) and gel purified. The vector and flanking regions were assembled using HiFi assembly mix (NEB) and transformed into electrocompetent PIR cells (ThermoFisher). Resulting plasmids were transformed into chemically competent MFDpir (66), which were used to generate merodiploids by conjugation with the wild-type strains. To select for the second crossover event and loss of the integrated plasmids, the cultures were spread on LB agar containing 20% sucrose and 0.5 ug/ml anhydrotetracycline for pFOK, or grown at 37°C on LB agar for pAX1. Successful gene deletions were identified using colony PCR targeting the gene of interest.

### Metal susceptibility tests

To measure growth in the presence of antimicrobial metals, a two-fold dilution series for CoCl2•6H_2_0 (Sigma Aldrich), CuSO_4_•6H_2_0 (Acros Organics), or anhydrous AgNO_3_ (Fisher Scientific) was prepared in Davis broth at concentrations ranging from 12.5 µM to 1 mM. An inoculum of approximately 2 x 10^6^ cells/ml was added to 2 ml Davis broth and incubated overnight. Growth in each sample was determined by spectrophotometry (OD600nm). To assess the growth with metal supplementation, stock solutions of FeCl_3_•6H_2_0 (Fisher Scientific) FeSO_4_•7H_2_0 (Fisher Scientific) and anhydrous MnSO_4_ were prepared in sterile water. The assays were performed as described above, using 800 µM Ga(NO_3_)_3_ concentrations in Davis broth without metal supplementation or supplemented with metals at concentrations ranging from 1.56 µM to 50 µM.

### β-galactosidase assay

The pTS12 vector was created by amplifying the kanamycin resistance gene and the ColE1 replication origin from plasmid pIDMv5K-J23100-fuGFP-B1006 (gifted from Sebastian Cocioba), and the promoterless *lacZ* gene from *E. coli* MG1655 genomic DNA (Q5 high fidelity mix, NEB). These two fragments were assembled using Gibson method with Hi-Fi DNA assembly master mix (NEB). The *fecA* promoter was inserted into pTS12 in front of *lacZ* by amplification from M12 genomic DNA followed by Gibson assembly with linearized pTS12. The pTS12 and *fecA::lacZ* reporter plasmids were verified by plasmid sequencing using Oxford Nanopore technology (Plasmidsaurus). The *fecA::lacZ* reporter plasmid and control pTS12 were transformed into strain M12 by electroporation, and β-galactosidase levels were measured by Miller assay as describe previously (67).

### Determination of intracellular iron, manganese, gallium, and zinc

Overnight cultures of M12 and M12 Δ*fecA* were subcultured in 25 ml of Davis broth with or without 200 µM Ga(NO_3_)_3_. Cultures were harvested after 4-5 hours and normalized to OD600 of 1.0. Cells were harvested by centrifugation, and washed twice in 25 ml 1X Tris-buffered saline, followed by a final wash in 20 mM Tris, 0.5 M sorbitol, and 200 uM EDTA (pH 7.5). Digestion of the final pellet was performed overnight in 1 ml of 68% concentrated nitric acid which was then diluted 10-fold in 2% nitric acid (prepared in chelex-treated milli-Q water) and analyzed by inductively coupled plasma mass spectrometry (ICP-MS) on an Agilent 7800 instrument. Calibration samples were made using each metal dissolved in nitric acid. Samples and standards were analyzed using the MassHunter software interface (Agilent).

## Statistical analyses

Analysis of variance (ANOVA) was carried out using Prism9 (GraphPad). A *P*-value <0.05 was considered statistically significant.

## Acknowledgements

This work was funded by the Marcus Jensen Research Endowment of Brigham Young University. The funders had no role in study design, data collection and interpretation, or the decision to submit the work for publication. We thank Adeyemi Ojaide for assistance with ICP-MS analysis and Sebastian Cocioba for providing plasmids.

